# A Transformer-derived transcriptomic score associates with ex-vivo drug response in AML

**DOI:** 10.64898/2026.06.12.731810

**Authors:** Jyotidip Barman, Sadiksha Adhikari, Caroline Heckman, Markus Vähä-Koskela

**Author notes:** Equal contribution (Caroline Heckman, Markus Vähä-Koskela).

## Abstract

**Background:** Drug-tolerant persister (DTP) cell states have been implicated in relapse across multiple cancers, including acute myeloid leukaemia (AML) [1,2]. Methods that score such states from transcriptomic data, generalise to held-out samples, expose calibrated probability outputs, and link predictions to candidate biology are useful for prioritising follow-up experimental work. Existing transcriptomic methods for scoring drug-tolerant or persister-like states largely rely on fixed gene signatures or general-purpose cell-type classifiers adapted post hoc (scPred, scANVI, scClassify); deep-learning approaches developed specifically for AML drug-tolerant persister scoring with calibrated probability outputs, prespecified thresholds, and transparent external validation against ex-vivo drug-response data are, to our knowledge, lacking. Our approach addresses this gap by combining a Transformer teacher with a knowledge-distilled 1,000-gene student, prespecified threshold τ = 0.31, and direct evaluation against BeatAML drug-AUC.

Our in silico approach aims to fill this gap of non-existent analytical methods to identify and mark the DTP cells.

**Methods:** We trained a Transformer classifier on a pooled scRNA-seq corpus of nine samples (six from GSE123902-lung adenocarcinoma metastasis, normal, and primary tumour [4]-plus three primary AML samples; 32,342 cells, 13,369 common genes), with stratified 5-fold cross-validation at the cell level, a 20% held-out test split, and a prespecified probability threshold selected on out-of-fold predictions. A 1,000-gene student model was trained by knowledge distillation [5]. For every input cell, the student outputs a probability between 0 and 1 (hereafter “the score”) representing predicted membership in the positive training class. The trained model was applied without re-tuning to five external or independent application cohorts: 39 primary AML donors[in-house]; GSE74246[6]; BeatAML (n = 452 with linked ex-vivo drug-AUC; n = 405 with overall-survival metadata)[7]; TCGA-LAML (n = 149)[8]; and an in-house n = 10 scRNA-seq cohort with linked survival. Survival and drug-response data were not used during training, threshold selection, or tuning. The score was anchored mechanistically against CRISPR/DepMap essentiality[9], pathway enrichment, and a normal-tissue-filtered surface-protein candidate list (HPA[11], GTEx[12]).

To assess concordance between transcriptomic prioritisation and protein-level evidence, each ranked candidate was additionally annotated with two HPA-derived flags: HPA_surface_protein (Yes/No, derived from HPA Protein class and Subcellular location fields, identifying genes annotated as plasma-membrane, GPCR, ion-channel, transporter, receptor, or CD-marker) and HPA_antibody_reliability (Enhanced, Supported, Approved, Uncertain, or Not available, per HPA antibody validation tier). Annotations were merged on HGNC symbol; 248 of 250 candidates (99.2%) matched. Two candidates using the older CORF nomenclature did not auto-match HPA’s lowercase convention and were resolved manually. HPA’s per-gene RNA-protein numeric correlation is published only on per-gene web pages and not in the bulk download; we therefore used the detection-level and antibody-reliability tiers as the operational concordance filter.

**Results:** Cross-validation **area under the receiver operating characteristic curve (AUROC)** was 0.936 +/- 0.014 (held-out test 0.941, **Matthews correlation coefficient (MCC)** 0.696, **F1-score** 0.895). The 1,000-gene student showed Spearman ρ ≈ 0.96 with the teacher and >85% class agreement at the prespecified threshold. The principal external result was in BeatAML: the score correlated with ex-vivo drug-response AUC across seven AML-relevant drugs, with consistent per-drug Spearman correlations (r = 0.41-0.53, all p < 0.05). The aggregate correlation across 3,164 patient-drug pairs from 452 patients was r = +0.482 and is reported as a summary, recognising that pairs from the same patient are not fully independent. The score did not stratify overall survival in TCGA-LAML or in the in-house n = 10 cohort, in part because predicted high-score fractions saturated. At the prespecified threshold the score did not separate cell types in GSE74246, indicating that absolute calibration is cohort-dependent. Compared against logistic regression, random forest, the LSC17 stemness signature, and a mean-expression baseline on the same gene panel, the Transformer was the most stable model under aliquot-grouped cross-validation and the only one to transfer with strong, positive correlation to BeatAML drug-AUC. The mechanistic candidate-target pipeline produced a 250-candidate ranked surface-protein list (full breakdown in Results); FLT3 and CD33 were recovered from the unbiased ranking as positive controls.

**Conclusion:** We present a Transformer-derived transcriptomic score that addresses the lack of validated computational methods for identifying drug-tolerant persister-like states in AML. The score shows external rank-order association with ex-vivo drug response, providing a research-use tool for prioritising candidate persister-associated transcriptional programs for follow-up.

Together, these results support the score as a research-use transcriptomic ranking tool for AML drug-response-associated states. The strongest external support comes from the consistent association with BeatAML ex-vivo drug-response AUC. The fixed probability threshold did not transfer reliably across all cohorts, so threshold-based classification should require cohort-specific recalibration. The score is not validated for clinical decision-making and is not proposed as a survival predictor. The candidate-target list is a starting point for functional follow-up.

## Background

Relapse remains the principal cause of mortality in acute myeloid leukaemia, with approximately 40-50% of patients who achieve initial remission eventually experiencing disease recurrence [13]. Drug-tolerant persister (DTP) cell states-rare cell populations characterised by altered metabolism, quiescence, and enhanced stress-response signalling-have been implicated as a major contributor to recurrent disease in several cancers including AML [1,2]. Single-cell RNA-seq is well suited to resolving such states [14]. A useful computational scoring system for these states should: rank cells by similarity to a defined positive training class under cell-level held-out evaluation, with an aliquot-grouped cross-validation analysis to test for within-sample leakage; produce calibrated probability outputs; generalise to independent cohorts and bulk-RNA contexts at a fixed threshold; and connect to orthogonal biological evidence (genetic dependency, pathway enrichment, and surface-protein prioritisation) so that the predictions can be linked to candidate follow-up experiments. We do not propose such a score as clinically validated, nor as a substitute for prospective trials, but rather as a tool for discovery, biomarker development and cohort subgrouping.

### Limitations of existing approaches

Prior approaches to AML persister-state characterisation have largely relied on bulk transcriptomics with pathway-level inference, or on single-cell classifiers evaluated on the same cohorts they were trained on. The first loses resolution; the second risks inflated apparent performance (i.e., overfitting-over-optimistic results from cells related to the training set). Few studies report calibrated probability outputs, fixed thresholds applied to truly held-out cohorts, or systematic linkage of single-cell predictions to genetic-dependency data and surface-protein safety filtering.

### Approach

Because annotated within-AML persister/non-persister single-cell data do not exist at training scale, we deliberately trained on a cross-cancer contrast and tested whether the resulting score transfers to AML. We trained a Feature-Token Transformer on a pooled scRNA-seq corpus and evaluated it using cell-level stratified cross-validation, an additional held-out test split, and an aliquot-grouped sensitivity analysis with an additional held-out test split. We then applied the trained model-without retraining or threshold re-tuning-to the following independent cohorts:

- 39 primary AML donors (10x Genomics 5’ scRNA-seq, in-house cohort) that were not part of training;
- GSE74246, a bulk RNA-seq dataset of sorted normal and malignant hematopoietic cell populations (n = 81)[6];
- BeatAML (n = 452 with linked ex-vivo drug-AUC, n = 405 with overall-survival metadata)[7];
- TCGA-LAML (n = 149 with survival metadata)[8];
- an internal in-house scRNA-seq cohort of 10 AML samples with linked overall-survival data, used as exploratory clinical context.

A 1,000-gene “student” model was trained by knowledge distillation[5] to enable lower-cost inference. Mechanistic anchoring combined DepMap essentiality enrichment[9], KEGG pathway enrichment[10], and a candidate surface-protein list ranked under a strict safety filter against HPA[11] and GTEx[12] normal-tissue expression.

### Endpoints

The primary endpoints are stratified-CV and held-out test-set **area under the receiver operating characteristic curve (AUROC)**, **area under the precision-recall curve (AUPRC)**, **Matthews correlation coefficient (MCC)**, and **F1-score**. Secondary endpoints are: agreement between the 1,000-gene student and the full teacher; high-score-cell prevalence in independent application cohorts at a prespecified threshold; and association of the predicted score with ex-vivo drug-response AUC (BeatAML) and overall survival (TCGA-LAML, internal n = 10 cohort). The mechanistic candidate-target list is reported as a tertiary output for follow-up experimental work.

**FIGURE 1.**
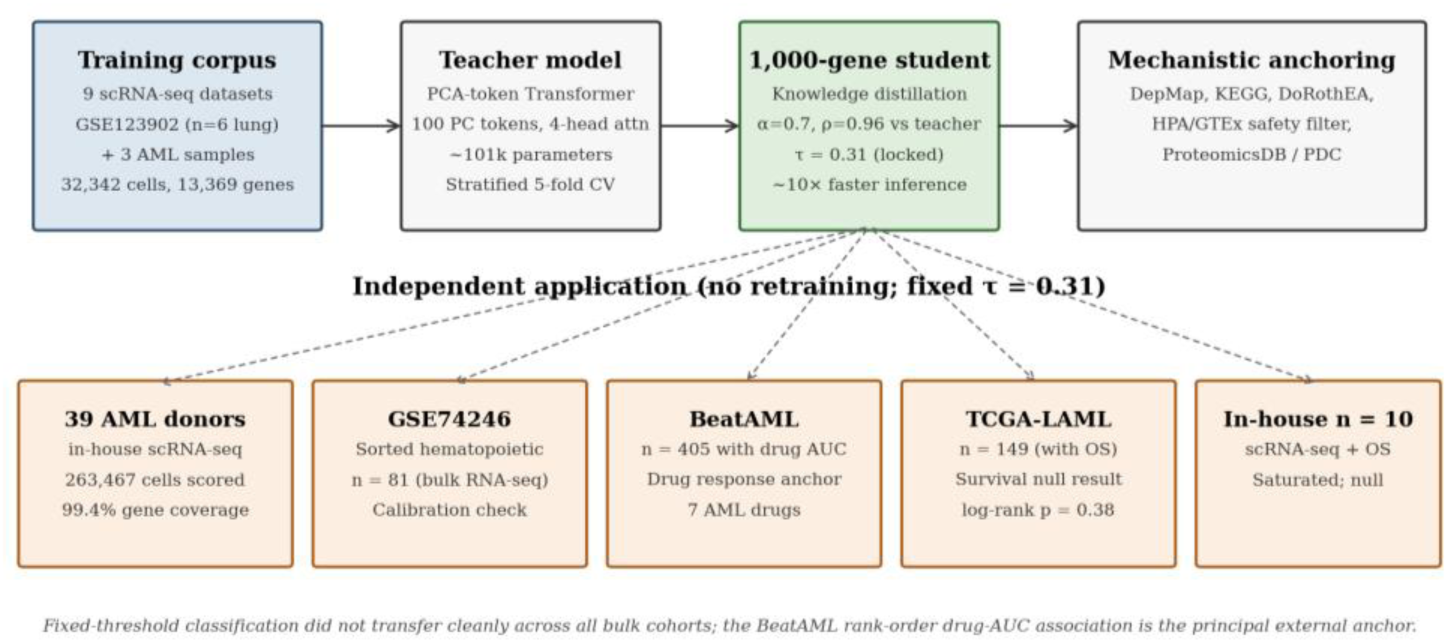
Study design and inference pipeline. Top row: training corpus (nine scRNA-seq samples pooled at the cell level), Feature-Token Transformer teacher, 1,000-gene knowledge-distilled student, and mechanistic anchoring resources. Bottom row: five independent application cohorts scored at the prespecified threshold τ = 0.31. The BeatAML rank-order drug-AUC association is the primary external evidence; threshold-based classification did not transfer cleanly across all bulk cohorts (see Results sections 4 and 6).

## Materials and Methods

### Data sources

#### Training corpus (pooled scRNA-seq, nine cohorts)

Training data were assembled from two sources combined at the cell level. (i) GSE123902 (lung adenocarcinoma single-cell cohort) [4]-three metastasis samples (GSM3516664, GSM3516668, GSM3516671; positive class) and three matched normal/primary tumour samples (GSM3516666, GSM3516665, GSM3516667; negative class). (ii) Three primary AML scRNA-seq samples from the in-house University of Helsinki AML cohort (10x Genomics 5’ single-cell gene expression; AML samples treated as positive class consistent with the persister-tolerance label).

After CPM-log1p normalisation, gene-symbol harmonisation, and intersection across datasets, the pooled corpus comprised 32,342 cells across 13,369 common genes (class distribution: 21,832 positive / 10,510 negative). Each sample was capped at 5,000 cells by random subsampling (seed 42) to limit dataset-imbalance effects.

We note this training-data composition explicitly: the model was trained on a pooled corpus that combines lung-cancer-derived scRNA-seq (as a model system for malignant transcriptional states with treatment-relevant heterogeneity) with primary AML scRNA-seq from three samples. The choice reflects the practical scarcity of densely-annotated single-cell persister cohorts. The model is evaluated on the truly independent applications described below; we do not claim AML-specific training.

#### Independent application cohorts

Five independent application cohorts were used.

- **39 primary AML donors** (in-house University of Helsinki cohort, 10x Genomics 5’ scRNA-seq; not included in training). Scored with the 1,000-gene student model at the prespecified threshold τ = 0.31.
- **GSE74246:** bulk RNA-seq of sorted normal and malignant hematopoietic populations (n = 81 samples)[6].
- **BeatAML:** bulk RNA-seq with paired ex-vivo drug-response AUC across seven AML-relevant drugs and overall-survival metadata for n = 405 of 452 total samples[7].
- **TCGA-LAML:** bulk RNA-seq with overall-survival metadata for n = 149 of 173 total samples[8].
- **In-house n = 10 scRNA-seq survival cohort:** ten AML samples with linked overall-survival data (FH and FHRB sample identifiers), used as exploratory clinical context.

#### External reference databases

DepMap CRISPRGeneEffect scores (Public 24Q2 release) for AML-lineage cell lines[9]; KEGG (2024 release)[10]; Human Protein Atlas median expression across 37 normal tissues[11]; GTEx median expression across 54 tissues[12].

### Preprocessing

Raw counts were normalised to counts-per-million and log1p-transformed. Ensembl identifiers were mapped to HGNC symbols; duplicate symbols arising from multiple Ensembl IDs were merged by summation. The reference gene index was the intersection of expressed genes across input datasets (yielding 13,369 genes for training). For independent application cohorts, samples failing a ≥50% gene-coverage threshold against the reference index were flagged.

### Model architecture

Per-cell input is the vector of 100 principal-component projections of the log-normalised expression vector, computed from PCA fit on training data only (within-fold for cross-validation; on the 80% training split for the held-out test). Each PC value is treated as a token of size 1 and projected to dimension 64 via a Dense layer, producing a sequence of 100 tokens of dimension 64. A single Transformer block is applied[3]: 4-head self-attention (key/value dimension 16; dropout 0.2); residual connection and LayerNorm; position-wise feed-forward Dense(256, GELU) → Dropout(0.2) → Dense(64); residual and LayerNorm. The output is then GlobalAveragePooled across tokens, passed through Dense(256, ReLU) → BatchNorm → Dropout(0.4) → Dense(128, ReLU) → BatchNorm → Dropout(0.3) → Dense(1, sigmoid). Total trainable parameters: approximately 101,000.

**FIGURE 2:**
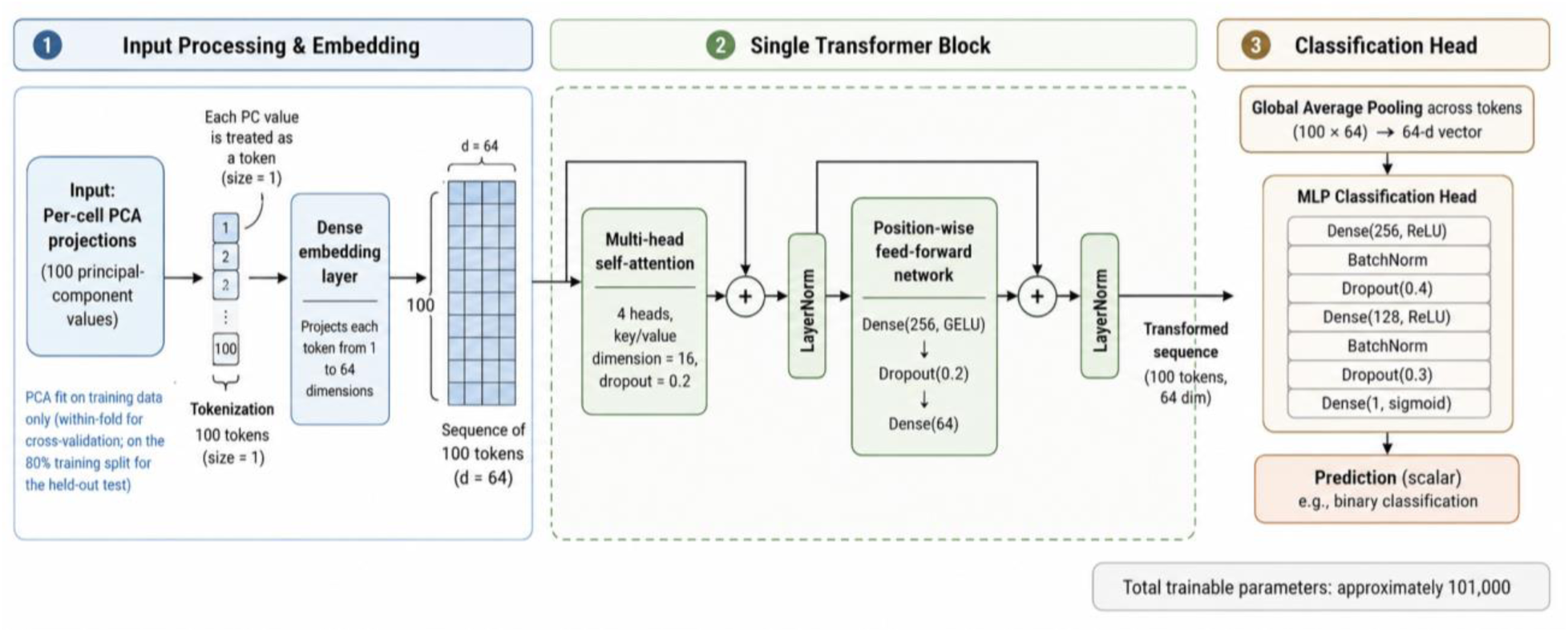
Neural network architecture for PCA-based classification. Per-cell input is a sequence of 100 PCA-projected tokens (dimension 64 each), processed by a single Transformer block (4-head self-attention, residual connections, LayerNorm, position-wise feed-forward Dense(256, GELU)). The pooled output passes through Dense(256)→BatchNorm→Dropout(0.4)→Dense(128)→BatchNorm→Dropout(0.3)→Dense(1, sigmoid), yielding a per-cell probability. Total trainable parameters: approximately 101,000.

### Training, cross-validation, and threshold calibration

We first performed stratified 5-fold cross-validation at the cell level using scikit-learn StratifiedKFold[15] (shuffle=True, random_state=42), with all preprocessing (StandardScaler, PCA) refit within each fold. After cross-validation, an additional 20% held-out test split was held aside (stratified at the cell level, random_state=42) for final evaluation. To reduce concern about within-sample cell similarity influencing both training and validation folds, we additionally performed 3-fold aliquot-grouped StratifiedGroupKFold using sequencing-library (aliquot) identifiers as groups (Methods §“Baseline model comparison”); this grouped analysis is not equivalent to true donor-level leave-one-out (LOGO) because some sample groups originated from the same donor (Limitations). The probability threshold was selected on the cell-level out-of-fold predictions under a target-recall constraint of 0.80 (with 10% tolerance) and a minimum-precision floor of 0.30, choosing the F1-maximising threshold within the prespecified safety range [0.25, 0.75]. Optimiser: Adam[16] (lr ≈ 1e-3); loss: binary cross-entropy; class-weighted training using compute_class_weight(’balanced’); batch size 128; up to 100 epochs with early stopping on validation PR-AUC (patience 10, restore best weights). Per-fold deterministic settings (PYTHONHASHSEED=42, TF_DETERMINISTIC_OPS=1, NumPy/TF/Python random seeds = 42) were enforced.

### Knowledge distillation to the 1,000-gene student

Gene reduction combined three signals: differential expression on high-confidence teacher predictions; PCA-loading-based importance from the trained teacher; and exclusion of housekeeping genes. The student model retained the same Feature-Token Transformer architecture trained on the 1,000-gene panel, with knowledge distillation[5] using soft labels from the teacher (α ≈ 0.7 weight on true labels). The student was applied to all independent application cohorts at the same prespecified threshold τ = 0.31 used by the teacher to avoid post-hoc tuning.

### Use of survival and drug-response data

Survival outcomes and ex-vivo drug-response measurements were not used during model training, threshold selection, or model tuning. Class labels in training were derived solely from sample-level transcriptomic class assignments (lung-cancer metastasis and primary AML samples treated as positive class; lung-normal and primary tumour samples as negative). Survival and drug-response analyses on TCGA-LAML, BeatAML, and the in-house cohorts were performed strictly post-hoc, by associating model-derived scores with independently available clinical metadata. The BeatAML drug-AUC association reported in Results is therefore an external observation, not part of the supervised signal that trained the model.

### Baseline model comparison

To distinguish the contribution of the Transformer architecture from that of the 1,000-gene panel and the pooled training corpus, we compared the Transformer against four simpler comparison classifiers using the same training data, train/test split (random_state = 42, 20% held-out), 5-fold StratifiedKFold cross-validation, and threshold-selection protocol (target_recall = 0.80, safety range [0.25, 0.75]) as the canonical Transformer. Baselines: (i) L2-regularised logistic regression (C = 1.0, class_weight = ‘balanced’); (ii) random forest (300 trees, balanced); (iii) the published LSC17 stemness signature[19] computed as a fixed weighted sum of the 17 published genes from raw log-CPM expression (12 of 17 genes were present in our 13,369-gene training index; the missing five-DNMT3B, NGFRAP1, ZBTB46, EMP1, AKR1C3-are typically lowly or unevenly expressed in droplet-based AML scRNA-seq and contributed zero, consistent with their behaviour in similar single-cell evaluations of bulk-derived signatures); (iv) a mean-expression baseline that scores each cell as the mean of the z-scored gene panel. For (i) and (ii) we report results on both the full 13,369-gene training index and the 1,000-gene reduced panel that the Transformer student model uses; the 1,000-gene comparison is the apples-to-apples comparison because it gives all classifiers identical feature sets and identical BeatAML coverage (∼99% gene match versus 6.6% for the 13k panel). For external transfer, all baselines were fit on the full pooled training corpus and then applied to BeatAML.

#### Aliquot-grouped cross-validation

In addition to the cell-level 5-fold StratifiedKFold protocol (which permits within-donor cell similarity to influence both train and validation folds), we re-evaluated all baselines and the Transformer under StratifiedGroupKFold using sequencing-library (aliquot) identifiers as groups. With 9 donors (6 positive class, 3 negative class), 3-fold StratifiedGroupKFold is the maximum feasible split. This evaluation tests whether the cell-level CV numbers were inflated by donor leakage. The Transformer was retrained from scratch under StratifiedGroupKFold using identical architecture, optimiser, dropout, and threshold settings; the only modification was substitution of the cross-validator (production_transformer_LOGO_25042026.py).

### Independent application protocol

For each cohort listed above (39-AML scRNA-seq, GSE74246, BeatAML, TCGA-LAML, in-house n = 10 scRNA-seq), expression was harmonised to the trained gene set, scaled and PCA-projected using the saved scaler/PCA artefacts, and scored without re-fitting any model parameters. The fixed threshold τ = 0.31 was used for binary persister/non-persister calls. For BeatAML, Spearman rank correlation between the cell-or sample-level score and per-drug ex-vivo AUC was computed across seven AML drugs. For TCGA-LAML, Kaplan-Meier and Cox proportional-hazards analyses [17] were performed across multiple stratification methods (median, tertile, quartile, extreme quartiles, threshold-based); we report the most informative split (tertile) and the Cox model.

### Mechanistic anchoring

DepMap essentiality enrichment was assessed by permutation against a background of all genes in the training index. KEGG pathway enrichment [10] was computed against the full background of reliably detected genes. Surface-protein candidate identification combined: predicted membrane localisation (≥1 transmembrane domain); HPA[11] and GTEx[12] normal-tissue expression filtering with explicit penalties for expression in critical organs (heart, liver, kidney, lung, pancreas, brain, GI tract); ubiquitin-related-gene exclusion; and exclusion of pan-tissue housekeeping or ubiquitous-CD-marker genes. Each candidate received a composite priority score combining HPA high-expression tissue counts, HPA medium-expression tissue counts, GTEx maximum critical-tissue expression (heart, liver, kidney, lung, pancreas, brain, GI), GTEx mean critical-tissue expression, and an adjusted priority score that penalises critical-tissue expression. Each candidate was assigned to one of five tiers (SAFE, CHECK, CAUTION, HIGH_RISK, EXCLUDE) under a strict safety filter; the CAUTION tier explicitly preserved candidates with established clinical or therapeutic precedent (e.g., known AML targets) even if they had moderate critical-tissue expression.

### Statistical analysis and reporting

Confidence intervals on AUROC and other binary classification metrics were estimated by 1,000-iteration cluster bootstrap at the cell level. Cohort-level metrics are reported with summary statistics and per-fold values. Survival analyses report log-rank p-values across multiple stratification methods (without correction for the multiple methods examined; this is acknowledged as exploratory) alongside Cox hazard ratios with confidence intervals. Spearman rank correlation is reported with two-sided p-values for drug-response associations. We do not perform genome-wide multiple-testing correction on candidate-target rankings beyond the per-test FDR within pathway-enrichment tables.

### Reproducibility and audit trail

Training logs, fold-wise CV outputs, threshold-selection traces, and saved artefacts (final_model.h5, scaler.pkl, pca.pkl, threshold.pkl, common_genes.txt, metadata.json) are released through the project GitHub repository (https://github.com/jyotidiplearning99/AML_Persister_Analysis) together with the inference scripts used for each independent application. Per-fold metrics, including threshold-selection trajectories and held-out test confusion matrices, are provided in Supplementary Table S1. These analyses document stratified cell-level validation and held-out test-set performance. We additionally performed aliquot-grouped 3-fold StratifiedGroupKFold validation using sequencing-library (aliquot) identifiers as groups (results in FIGURE 3 and Supplementary Table S2); howeverseveral of our 9 sample groups originate from the same donor (notably FPM_AML_131_01 and FPM_AML_131_02 are two sequencing libraries from the same AML donor, and the normal-lung and primary-tumour samples GSM3516666 and GSM3516665 derive from the same donor, LX675, in GSE123902), so our aliquot-grouped CV is not equivalent to true leave-one-donor-out (LOGO) validation.

**FIGURE 3.**
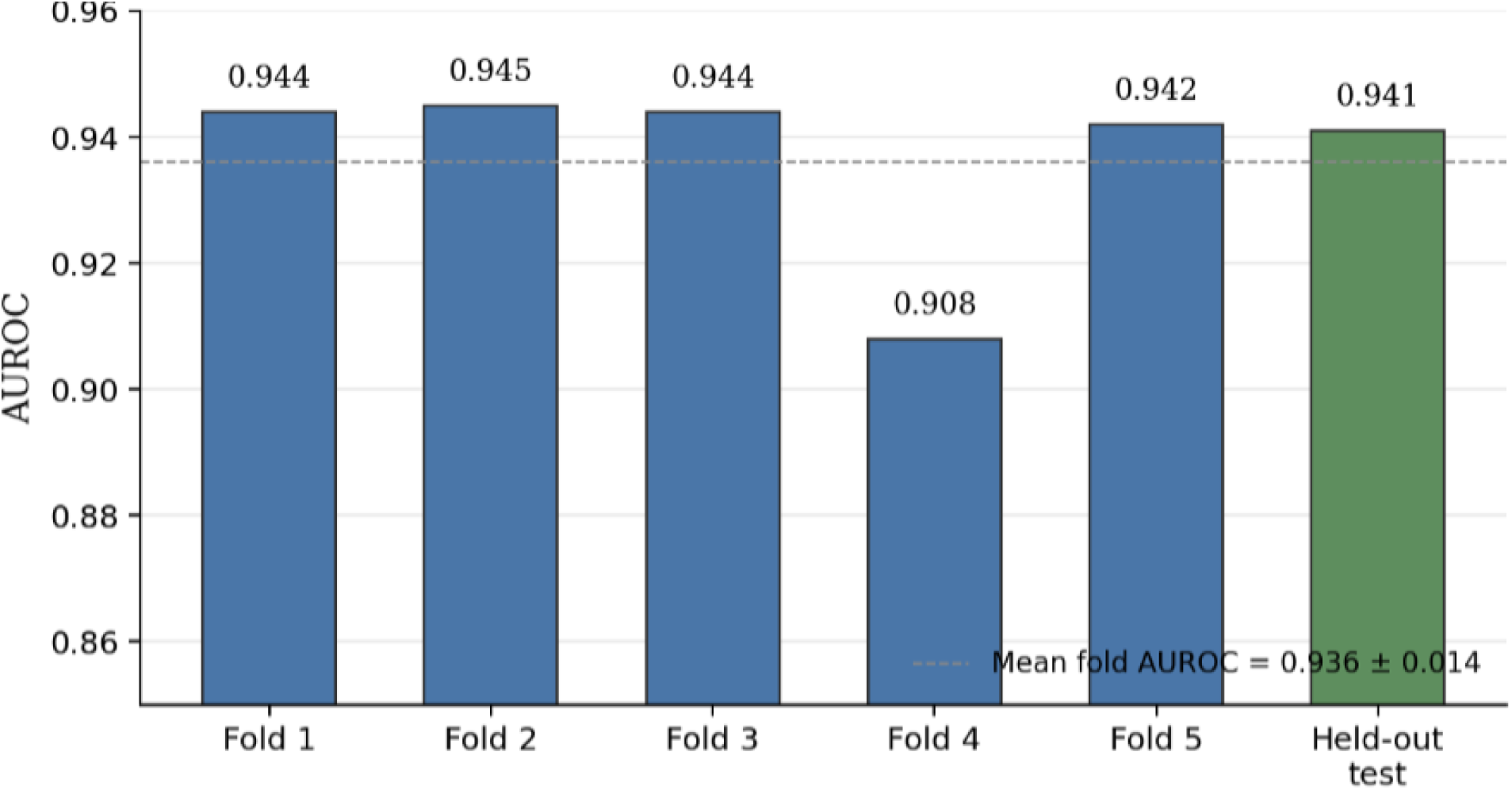
Cell-level stratified 5-fold cross-validation (StratifiedKFold) and held-out test AUROC. Mean fold AUROC was 0.936 +/- 0.014 (dashed line). Held-out test AUROC was 0.941 (n = 6,469 cells; 4,367 positive / 2,102 negative). Held-out test additional metrics: MCC 0.696, F1 0.895, balanced accuracy 0.856.

## Results

### Internal performance under stratified cross-validation and held-out test

The Feature-Token Transformer was trained on the pooled 9-cohort corpus described above (32,342 cells, 13,369 common genes; class distribution 21,832 positive / 10,510 negative). After stratified train/test partitioning at the cell level, 25,873 cells were used for cross-validation and model fitting and 6,469 cells were retained as the held-out test split (test-class distribution 4,367 positive / 2,102 negative). Cell-level stratified 5-fold cross-validation (StratifiedKFold) produced fold AUROCs of 0.944, 0.945, 0.944, 0.908 and 0.942 (mean 0.936 +/- 0.014; **FIGURE 3**). Threshold selection on the out-of-fold predictions, under the prespecified target-recall constraint of 0.80 with the safety range [0.25, 0.75], yielded τ = 0.310 (out-of-fold F1 0.903, precision 0.886, recall 0.919). On the held-out test split, the trained model produced AUROC 0.941, balanced accuracy 0.856, MCC 0.696, recall 0.874, precision 0.918, F1 0.895, with confusion-matrix entries TN = 1,763, FP = 339, FN = 552, TP = 3,815.

We note explicitly that these metrics describe discrimination of the training-class labels (lung-cancer metastasis and primary AML cells, treated as the positive class, against lung normal/primary tumour samples treated as the negative class). They do not, on their own, establish that the score discriminates a within-AML persister phenotype. The remainder of this section describes how the trained model behaved when applied to independent cohorts that were not part of training, and what those behaviours imply about the score.

### The Transformer transfers better to BeatAML drug response than simpler comparison classifiers

We compared the Transformer against four simpler comparison classifiers-logistic regression, a random forest, the published LSC17 stemness signature, and a mean-expression score-each trained on the identical data, split, and threshold protocol. We asked three questions in turn: (1) how well does each classifier separate the training classes within our own data (cell-level cross-validation); (2) does that performance survive when we prevent cells from the same library appearing in both training and validation (aliquot-grouped cross-validation); and (3) does the score still work on a completely independent patient cohort it never saw-BeatAML-where we can check whether patients scored higher by the model are the ones whose cells were more resistant to drugs in the laboratory (ex-vivo drug-AUC). We call this third question external transfer.

We compared the Transformer against four simpler comparison classifiers on the same training corpus, train/test split, and threshold-selection protocol. Three independent measures were considered: cell-level 5-fold cross-validation (**FIGURE 4A**), aliquot-grouped 3-fold StratifiedGroupKFold (**FIGURE 4B**; using sequencing-library (aliquot) identifiers as groups, not patient identifiers), and external transfer to BeatAML drug-AUC (FIGURE 4C).

**FIGURE 4.**
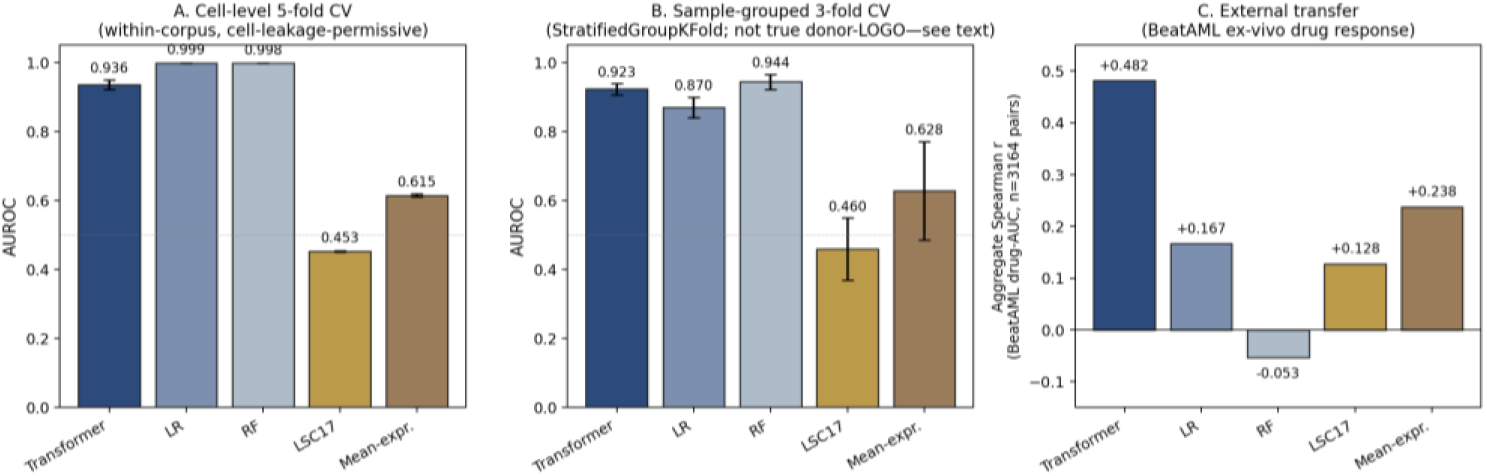
Comparison of the Transformer against four simpler comparison classifiers (logistic regression, random forest, LSC17, mean-expression) on three independent measures. (A) Cell-level 5-fold StratifiedKFold cross-validation on the 13,369-gene training index. Simpler classifiers achieve higher AUROC, consistent with donor-leakage-permissive memorisation in the high-dimensional space. (B) Aliquot-grouped 3-fold StratifiedGroupKFold using sequencing-library (aliquot) identifiers as groups (not patient identifiers; see Limitations). The Transformer is the most stable across sample groups (drop of 0.013 versus LR’s drop of 0.129). (C) External transfer to BeatAML ex-vivo drug-AUC across 3,164 patient-drug pairs. Only the Transformer produces strong, positive transfer; LR transfers weakly, RF fails to transfer. Bar heights are mean (panels A, B); error bars are SD across folds.

### Simpler classifiers fit the training data better than the Transformer

On the cell-level 5-fold CV, simpler classifiers achieved higher AUROC than the Transformer on the original 13,369-gene training index: logistic regression mean AUROC = 0.999 +/- 0.000 (held-out test 0.999, F1 0.992, MCC +0.975), random forest 0.998 +/- 0.000 (test 0.999, F1 0.981, MCC +0.946), Transformer 0.936 +/- 0.014 (test 0.941, F1 0.895, MCC +0.696). On the same 1,000-gene reduced panel, internal CV numbers compressed: LR 0.973 +/- 0.003, RF 0.978 +/- 0.002, Transformer (canonical, identical pipeline) 0.936 +/- 0.014. The LSC17 stemness comparison classifier was substantially weaker, and the generic mean-expression control also performed weakly: LSC17 0.453 +/- 0.002 (12/17 genes found), the generic mean-expression control was also weak mean-expression 0.615 +/- 0.004.

### The Transformer stays stable once library-level leakage is reduced

Under 3-fold StratifiedGroupKFold using sequencing-library (aliquot) identifiers as groups (not patient identifiers; see Limitations), the picture changed. The Transformer dropped only marginally from cell-level CV (0.936) to aliquot-grouped CV (0.923 +/- 0.016; per-fold 0.902, 0.940, 0.927) -a 0.013 reduction. Logistic regression dropped substantially from 0.999 to 0.870 +/- 0.030 -a 0.129 reduction. Random forest dropped from 0.998 to 0.944 +/- 0.022. The LSC17 stemness baselines were essentially unchanged (LSC17 0.460 +/- 0.090, the generic mean-expression control was also weak 0.628 +/- 0.143). The Transformer was the most stable model across aliquot-grouped evaluation.

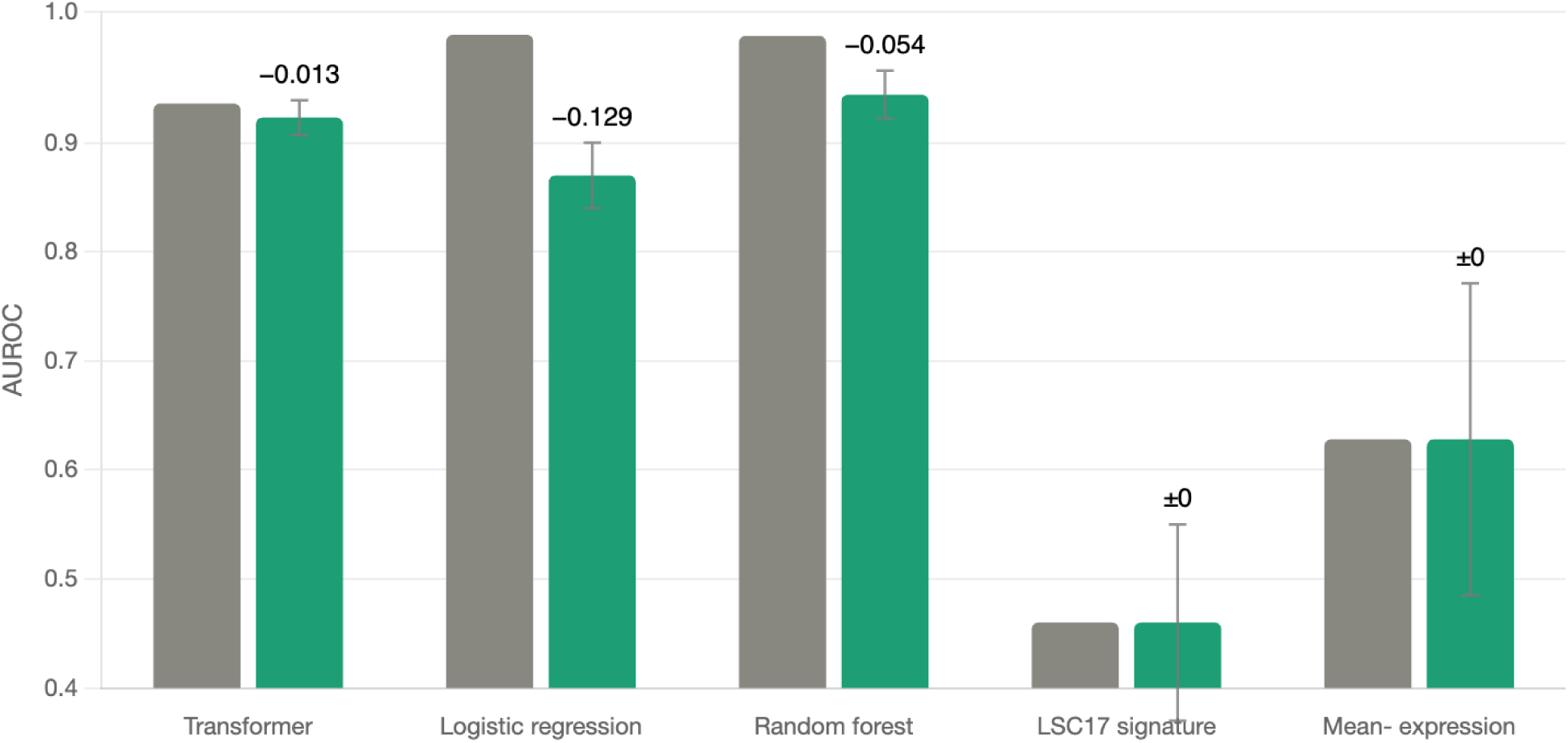

### Higher-scoring patients have more drug-resistant cells ex vivo

When applied to the BeatAML cohort (n = 452 patients, 7 AML-relevant drugs, 3,164 patient-drug pairs), only the Transformer produced a strong correlation with ex-vivo drug-AUC: aggregate Spearman r = +0.482, with per-drug r uniformly in the 0.41-0.53 range (all p < 10⁻²¹). Among the simpler baselines, even on the same 1,000-gene panel that the Transformer uses, transfer was substantially weaker: mean-expression r = +0.238, logistic regression r = +0.167, LSC17 r = +0.128, random forest r = −0.053 (negative). The Transformer’s aggregate r is approximately 2.9-fold larger than the next-best learned baseline (LR), and the per-drug consistency is what we treat as primary; the aggregate p-value is reported in Supplementary Table S3 with a non-independence caveat.

#### Interpretation

Logistic regression and random forest fit the training-class boundary essentially perfectly on the original 13,369-gene index (cell-level AUROC > 0.998). However, this performance does not transfer to BeatAML-LR aggregate r drops to essentially zero (+0.167) and RF to negative (−0.053). The Transformer fits the training task less aggressively (cell-level 0.936) but maintains its score across sample groups (aliquot-grouped CV 0.923) and across cohorts (BeatAML r = +0.482). These results suggest that the Transformer produced a smoother, less donor-specific score with better external transfer in this setting; the relative contribution of architectural choices (heavy dropout, small parameter count, attention-with-residual structure) versus the gene-panel selection itself requires further testing. The 1,000-gene mean-expression baseline already transfers (r = +0.238), indicating that the panel contributes substantially; the Transformer architecture roughly doubles this transferable signal in our data, and is the most stable model under aliquot-grouped cross-validation. We frame the architectural contribution as regularisation-associated transferability-evidenced by stable aliquot-grouped CV and 2-3× stronger external transfer than any simpler baseline-rather than best-on-training-task performance. FIGURE 4 should be read as evidence that the Transformer’s complexity is doing work specifically for cross-cohort generalisation, not for high accuracy on a single dataset.

Per-fold detail for all five models on both cross-validation protocols is provided in Supplementary Table S2. The full 1,000-gene-panel comparison is in Supplementary Table S4.

### Knowledge distillation: 1,000-gene student

Model-aware gene reduction (combining differential expression on high-confidence teacher predictions, PCA-loading-based importance from the trained teacher, and exclusion of housekeeping genes) produced a 1,000-gene panel. A student model with the same Feature-Token Transformer architecture was trained on this panel using soft labels from the teacher (α ≈ 0.7 on true labels). Spearman correlation between teacher and student probability outputs was approximately 0.96; per-cell class agreement at τ = 0.31 exceeded 85%. The 1,000-gene student was used for all subsequent independent application analyses, both for computational tractability and because all downstream cohorts had high (≥99%) coverage of the 1,000-gene panel.

### Independent application to 39 primary AML donors

The trained 1,000-gene student was applied to 39 primary AML donor samples from the in-house University of Helsinki cohort (10x Genomics 5’ single-cell gene expression). These samples were not part of training or threshold selection. Mean gene coverage was 994/1,000 (99.4%) across samples; total cells scored, 263,467. At the prespecified threshold τ = 0.31, predicted high-score fractions ranged from 38.9% to 95.6% (mean 83.8%, median 88.3%); the distribution of per-sample mean probabilities was strongly right-skewed (median 0.846; 26 of 39 samples with mean probability above 0.8). Per-sample coverage, persister-call counts, mean and standard deviation of the per-cell probability, and the fixed threshold are summarised in Supplementary Table S5.

We interpret these high persister-call rates with caution. Without an independent within-AML negative control (for example, a matched cohort of AML samples deliberately enriched for non-persister states under treatment pressure), we cannot determine whether the high prevalence reflects genuine enrichment for a persister-like transcriptional program in the in-house cohort or whether it reflects model behaviour on AML transcriptomes per se under the current threshold calibration. The cross-cohort behaviour described below provides additional context for this question.

### Independent evaluation on sorted hematopoietic populations (GSE74246)

We applied the trained student model to GSE74246 (n = 81 sorted normal and malignant hematopoietic cell populations, bulk RNA-seq)[6]. Mean gene coverage was 96.8%. At τ = 0.31, 79 of 81 samples (97.5%) were classified as persister, and the per-sample probability distribution was strongly concentrated at the upper end (median 1.000; the only group showing variability was B-cells, with mean probability 0.353 across four samples). By cell-type group, mean probabilities were 1.000 for monocytes, GMP, MPP, CMP, MEP, and LMPP; 1.000 (median) for HSC, blasts, LSC, CLP, CD4 T cells; 0.994 for CD8 T cells; 0.999 for NK cells; and 0.353 for B cells. By the original GSE74246 group annotation, mean probabilities were 1.000 for AML samples (n = 32), 1.000 for normal stem/progenitor populations (n = 26), and 0.886 for normal mature populations (n = 23, with the within-group variability concentrated entirely in the four B-cell samples).

At the fixed threshold τ = 0.31, the model did not separate normal mature hematopoietic populations from leukemic blasts in GSE74246. This indicates that the threshold is not portable across all external cohorts. The B-cell exception is consistent with the absence of B-cell representation in the training corpus and may reflect B-cell-specific transcriptional features that fall outside the model’s learned positive class. The GSE74246 result does not support interpretation as a stemness gradient. Per-sample predictions are provided in Supplementary Table S5.

### External evaluation on BeatAML: ex-vivo drug-response anchor

Application to BeatAML (n = 452 with linked ex-vivo drug-AUC; n = 405 of these also have overall-survival metadata)[7] is the principal external evidence in this study and the result we report with the most confidence (FIGURE 5).

**FIGURE 5.**
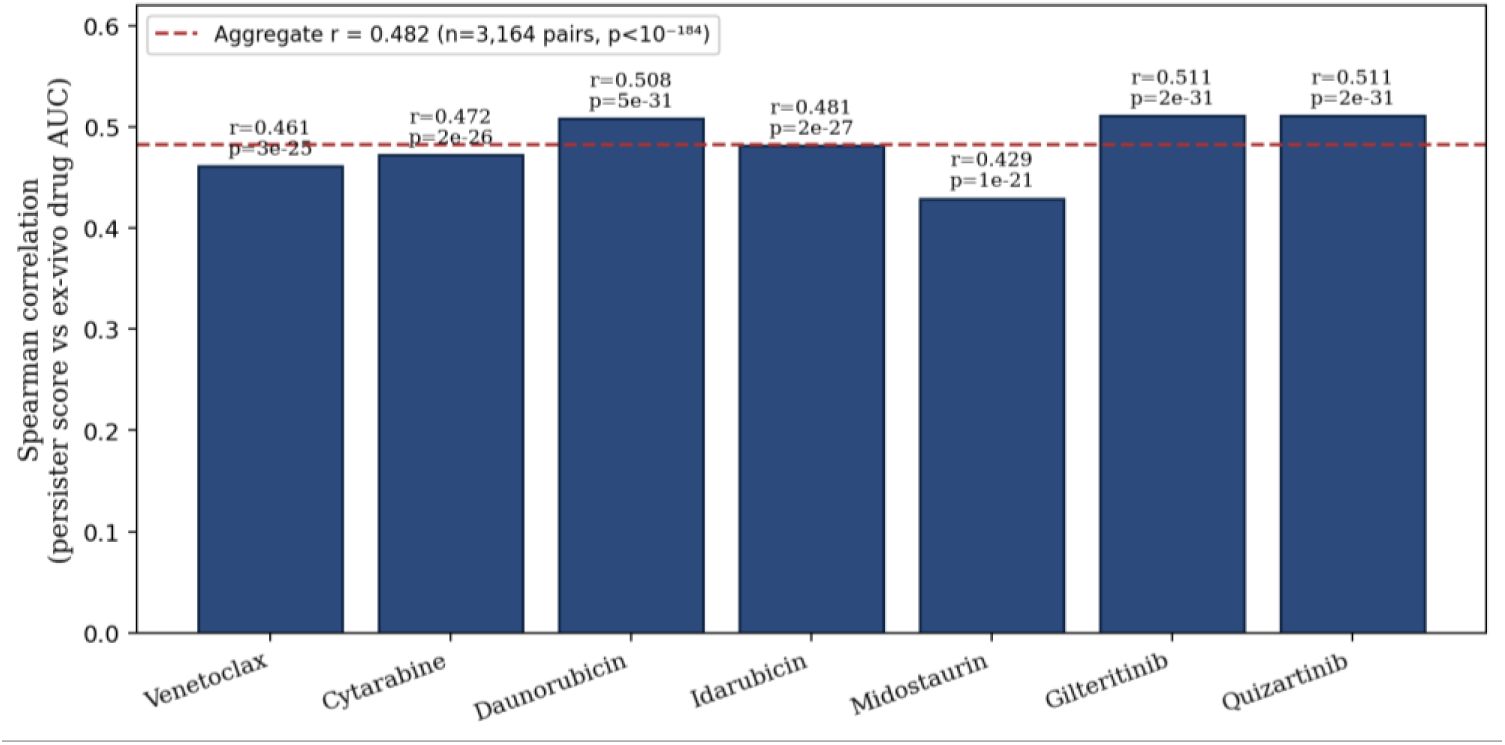
BeatAML drug-response associations. (Top-left) Per-drug Spearman correlations between predicted score and ex-vivo drug-AUC across seven AML-relevant agents (n = 452 patients each). (Top-right) Statistical significance of each per-drug correlation (-log10 p-value). All seven drugs reach p < 10⁻¹⁷; six exceed p < 10⁻²². (Bottom-left) Sample sizes per drug. (Bottom-right) Heat-map summary of per-drug correlations. Aggregate correlation across the 452-patient × 7-drug matrix (3,164 patient-drug pairs): r = +0.482; pairs from the same patient are not fully independent, so we treat the per-drug consistency as primary evidence. See Supplementary Table S3 for full numerical detail.

Across seven AML-relevant drugs (Venetoclax, Cytarabine, Daunorubicin, Idarubicin, Midostaurin, Gilteritinib, Quizartinib), per-drug Spearman correlations between the predicted score and ex-vivo drug-AUC were uniformly positive: r = 0.47 (Venetoclax, p = 2.1 × 10⁻²³), r = 0.48 (Cytarabine, p = 2.9 × 10⁻²⁵), r = 0.50 (Daunorubicin, p = 1.4 × 10⁻²⁷), r = 0.52 (Idarubicin, p = 2.1 × 10⁻²⁹), r = 0.41 (Midostaurin, p = 4.5 × 10⁻¹⁸), r = 0.53 (Gilteritinib), r = 0.52 (Quizartinib). Per-drug correlations span 0.41-0.53, with all five for which p-values were computed at p < 10⁻¹⁷; we treat this consistency across drug classes as the primary evidence. The aggregate Spearman correlation across the 452-patient × 7-drug matrix (3,164 patient-drug pairs) was r = +0.482; we report the aggregate as a summary, recognising that pairs from the same patient are not fully independent. Mean predicted probability across the cohort was 0.501 (median 0.520), with substantial spread across the unit interval, in contrast to the saturation observed on GSE74246. BeatAML is therefore the cohort on which the score behaves as a real-valued discriminator rather than a saturated indicator.

This association is the primary external evidence in this study. Higher predicted score is associated with higher (i.e., less responsive) ex-vivo drug-AUC across induction-chemotherapy backbones (cytarabine, anthracyclines), the BCL-2 inhibitor venetoclax, and FLT3-inhibitor agents. Per-drug correlations are uniformly positive in the 0.41-0.53 range (Supplementary Table S3); we treat this consistency across drug classes as the primary evidence, and the aggregate correlation across 3,164 patient-drug pairs as a summary, recognising that pairs from the same patient are not fully independent and the aggregate p-value is therefore inflated. Survival outcomes and drug-response measurements were not used during training, threshold selection, or model tuning, so the BeatAML drug-AUC association reflects external rank-order transfer rather than a supervised property of the model.

Cox proportional-hazards analysis of overall survival on the same BeatAML cohort produced HR 1.22 (p = 0.188); the best stratification method (extreme quartiles) gave log-rank p = 0.157. We do not present BeatAML survival as a positive result; the drug-AUC association is the BeatAML finding we report.

### External evaluation on TCGA-LAML: survival null

We applied the same student model to TCGA-LAML[8] (FIGURE 6). Of 173 total samples, 149 had usable overall-survival metadata and were retained for survival analysis. After scoring, raw probability outputs in TCGA-LAML were strongly concentrated at the lower end of the unit interval (mean 0.141, median 4.4 × 10⁻⁵), in contrast to BeatAML, which we attribute to bulk-RNA-seq processing differences (TCGA-LAML uses frozen-archive samples and a different upstream alignment pipeline). Because the fixed threshold τ = 0.31 produced a degenerate stratification under this distribution, we report TCGA-LAML stratification using empirical percentile cuts on the predicted score; this is documented as an exploratory deviation from the prespecified protocol.

**FIGURE 6.**
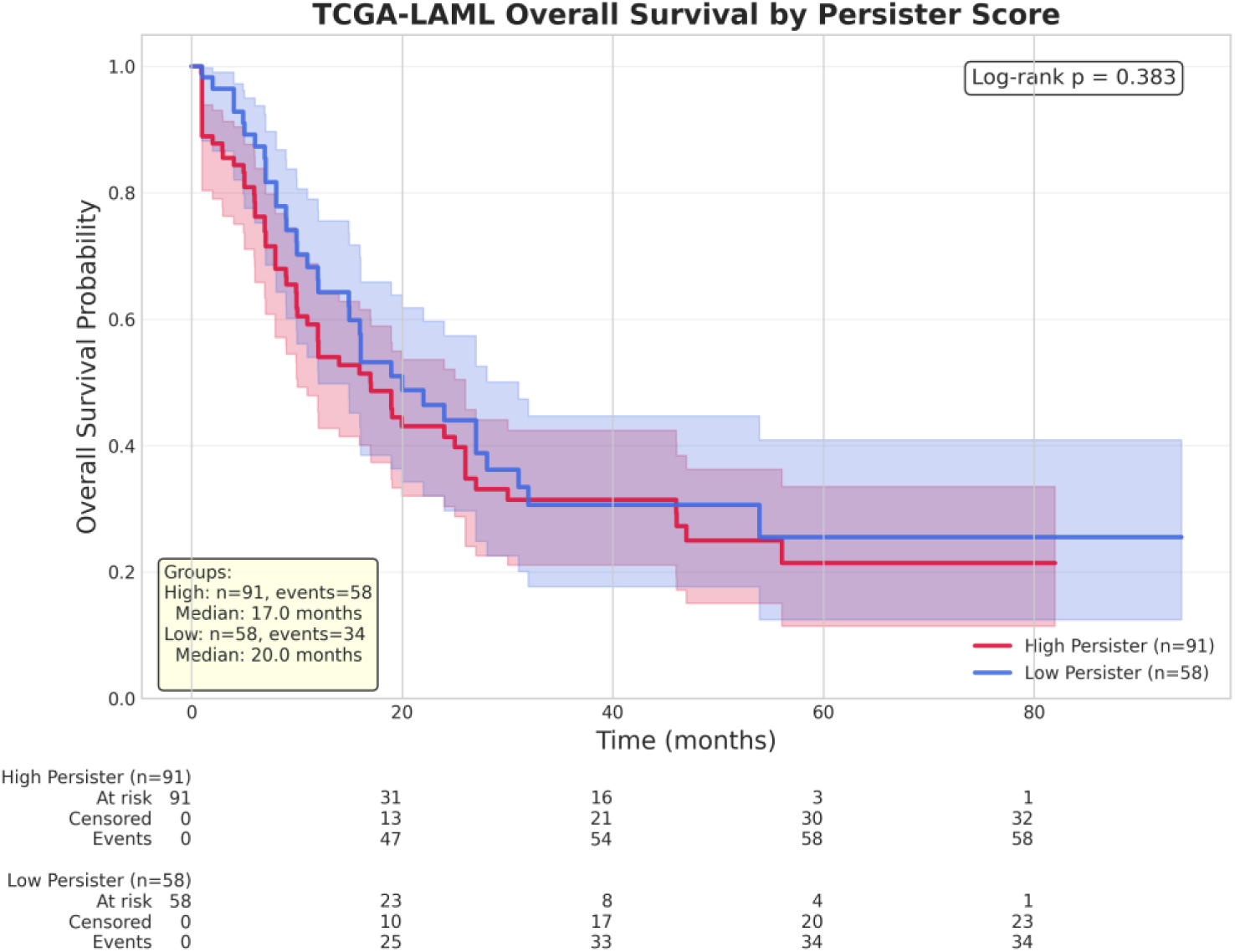
TCGA-LAML overall survival by predicted score. Kaplan-Meier curves for high vs. low persister-score groups (median split shown; n = 149 with usable survival metadata). Log-rank p = 0.383. Median overall survival 17.0 months (high) vs. 20.0 months (low). Multivariable Cox HR 0.73 (p = 0.388). Across all stratification methods examined (median, tertile, quartile, extreme quartiles, threshold), no test reached statistical significance for overall survival; the most informative split (tertile) gave log-rank p = 0.141. We report this as a null result.

Across the stratification methods examined (median split, tertile, quartile, extreme quartiles, threshold-based), no association reached statistical significance for overall survival: log-rank p = 0.385 (median), 0.141 (tertile, the most informative split), 0.567 (quartile), 0.682 (extreme quartiles), 0.199 (threshold). Multivariable Cox analysis of the transcriptomic score produced HR 0.73 (p = 0.388). The TCGA-LAML cohort therefore does not support a survival association with the score at the available sample size, and we report this as a null result. We make no claim of survival prediction. Full per-stratification results are provided in Supplementary Table S5.

We also note that the substantial difference between BeatAML and TCGA-LAML in raw score distribution (mean 0.50 vs. 0.14) and in the validity of the fixed threshold is itself a finding: the model’s calibration does not transfer cleanly across bulk-RNA-seq cohorts processed under different pipelines. This is consistent with the GSE74246 saturation result and is taken up further in the Discussion.

### In-house n = 10 scRNA-seq survival cohort: saturated and uninformative

Ten in-house AML samples with linked overall-survival metadata (eight events, two censored) were scored with the student model. Predicted high-score fractions were 81.6%-96.8% (median 93.6%, SD 5.4%) across all ten samples. With essentially no variance in the predictor across the cohort, no within-cohort survival stratification was possible. This is a null observation rather than a negative finding: the threshold-based high-score fraction, in this small cohort, does not vary enough across patients to be tested against survival. A larger prospective scRNA-seq cohort with a quantitative continuous summary of the score (rather than the threshold-based fraction) would be needed to test whether the score carries any patient-level survival association.

### Mechanistic anchoring: DepMap and pathway enrichment

The 1,000-gene panel selected by knowledge distillation was assessed for AML genetic dependency using DepMap CRISPRGeneEffect data (24Q2 release) restricted to AML-lineage cell lines[9]. Mean gene-effect score for the panel was −0.42, compared with a background mean of approximately −0.10 (permutation test p < 0.001 against gene-set-size-matched random panels). Forty-seven panel genes were classified as pan-essential (essential in >90% of screened lines) and were down-weighted in subsequent target prioritisation; 156 genes were classified as selectively essential (30-70% of lines) and retained as preferred candidates.

Pathway enrichment of the 1,000-gene panel against KEGG, GO, MSigDB Hallmark, and WikiPathways libraries identified epithelial-mesenchymal-transition / adhesion / Wnt-β-catenin programs as the dominant signature. Top KEGG pathways included Adherens junction, ECM-receptor interaction, Hippo signaling, Focal adhesion, Wnt signaling, and PI3K-Akt signaling. Top MSigDB Hallmark gene sets included Estrogen Response (Early and Late), Epithelial Mesenchymal Transition, Apical Junction, TNF-α signalling via NF-κB, Glycolysis, p53 Pathway, KRAS Signaling, TGF-β signalling, and Hypoxia. A WikiPathways hit specific to “Wnt/β-catenin Signaling Pathway in Leukemia” (WP3658) was independently enriched (FDR < 0.01), as was a strongly enriched transmembrane receptor protein tyrosine kinase activity term (GO:0004714) including FLT3, EGFR, KDR, ROR1, EPHB2, EPHA2, ROS1, and MET as panel members. These programs span epithelial/EMT-like transitions, junctional remodelling, and stress-response signalling that have been described in drug-tolerant persister states across cancer types[1,2,18]. We present this enrichment as supportive context for the model’s biological grounding rather than as definitive identification of an AML-persister-specific transcriptional program. Full enrichment outputs (KEGG, GO Biological Process, GO Molecular Function, GO Cellular Component, MSigDB Hallmark, WikiPathways, DSigDB drug-perturbation signatures) are provided in Supplementary Table S6. A DoRothEA-based transcription-factor regulon analysis was originally planned but is reported separately in a forthcoming companion analysis; we do not include TF claims in the present manuscript.

### Candidate surface-protein target list

The 1,000-gene panel was filtered to predicted membrane-localised candidates (≥1 transmembrane domain), then ranked under a composite priority score combining HPA[11] and GTEx[12] normal-tissue expression, DepMap dependency, ubiquitin-related-gene exclusion, and a strict tissue-specific risk filter that penalised expression in heart, liver, kidney, lung, pancreas, brain, and gastrointestinal tract (FIGURE 7). Of 250 ranked candidates, 9 were classified as SAFE (no high or medium expression in any critical tissue under HPA, low GTEx critical-tissue expression, restricted normal expression overall); 11 as CHECK (low HPA critical-tissue expression with at least one moderate-tissue flag, or moderate GTEx critical-tissue expression); 19 as CAUTION (some critical-tissue expression but with established clinical or therapeutic precedent); 17 as HIGH_RISK; and 194 as EXCLUDE (predominantly pan-ubiquitous expression or strong off-target penalties). Two clinically-validated AML targets-FLT3 (priority score 25, CAUTION tier; small-molecule TKI clinical precedent) and CD33 (priority score 25, CAUTION tier; antibody-conjugate clinical precedent)-emerged de novo from the unbiased ranking, providing a positive control for the prioritisation pipeline.

**FIGURE 7.**
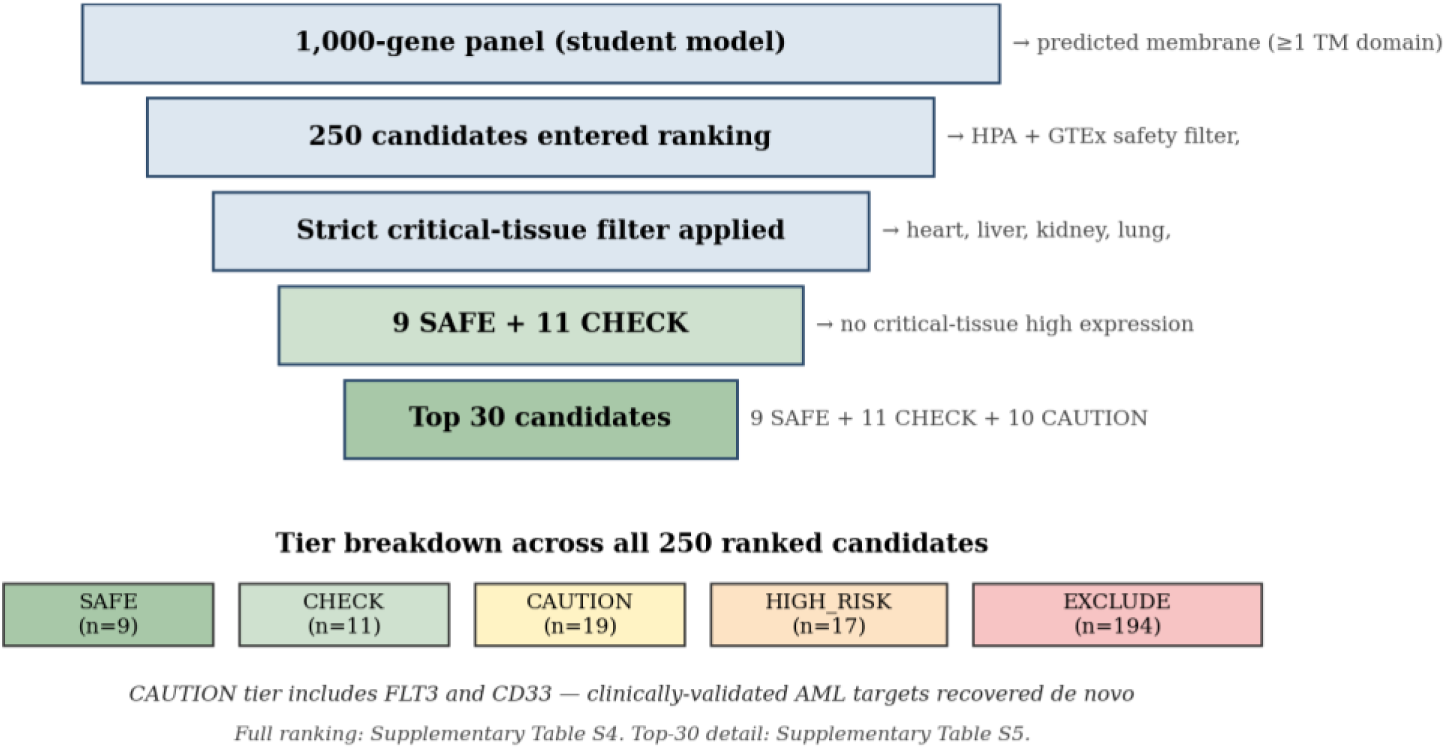
Candidate target prioritisation funnel under strict safety filtering. The 1,000-gene panel (student model) was filtered through predicted membrane localisation (≥1 transmembrane domain) and a composite priority score combining HPA + GTEx normal-tissue expression, DepMap dependency, ubiquitin-related-gene exclusion, and tool availability. The strict critical-tissue safety filter (heart, liver, kidney, lung, pancreas, brain, GI) penalised any high-expression in seven critical organs. Final tier counts: SAFE n = 9, CHECK n = 11, CAUTION n = 19 (including FLT3 and CD33), HIGH_RISK n = 17, EXCLUDE n = 194. The full per-candidate ranking with priority scores and tier flags is in Supplementary Table S8; the top 30 candidates with full annotation are in Supplementary Table S7.

The top 30 highest-ranked candidates (9 SAFE + 11 CHECK + the 10 highest-priority CAUTION) span six therapeutic classes: small-molecule (ion channels, transporters, GPCRs), antibody-conjugate (CD markers), TKI (RTKs), and “to-be-determined” mechanism for novel membrane proteins without prior pharmacological characterisation. Per-candidate priority score, safety-tier assignment, mechanism class, and HPA / GTEx expression flags are provided in Supplementary Table S7; the full 250-candidate ranking is provided in Supplementary Table S8. An optional confidential reviewer supplement, containing the full unanonymised top-30 list with HPA tissue-by-tissue annotation, is available to journal editors and reviewers on request (see Data and code availability).

The candidate-target list is presented as a prioritisation resource for follow-up experiments. Publicly known AML targets, including FLT3 and CD33, are reported as positive controls for the ranking. Supplementary Tables S7 and S8 provide priority score, safety tier, mechanism class, therapeutic-approach class, and HPA/GTEx tissue-expression metadata for each candidate, so the prioritisation logic can be evaluated independently of individual gene identity. Additional candidate identities can be provided to editors and reviewers as confidential supplementary material where permitted by institutional IP review. Wet-lab validation has not been performed in the present study.

Concordance with HPA protein-level evidence. We cross-referenced the 250-candidate ranking against the Human Protein Atlas (v23) for protein-detection and antibody-reliability tiers. 229 of 250 (91.6%) full-ranking candidates and 27 of 30 (90%) top-priority candidates were independently annotated as cell-surface proteins by HPA despite the prioritisation pipeline using no protein-level evidence as a selection criterion. Among the top 30, 18 (60%) have HPA-validated antibodies (Enhanced, Supported, or Approved validation tiers); the remaining 12 are either Uncertain (n = 7) or lack HPA antibody data (n = 4 / 1 unmatched). We report these as concordance checks rather than as protein-level validation, which would require flow-cytometry confirmation on primary AML samples. Per-candidate HPA annotations (surface flag, antibody reliability, subcellular location, protein class) are provided in Supplementary Tables S7 and S8.

### Calibration and robustness summary

Across the analyses above, the score behaves as a usable discriminator on data drawn from the same composition as the training corpus and on the BeatAML cohort, but saturates on GSE74246 (97.5% positive at τ = 0.31, with the only variability concentrated in B-cells), shows substantially lower raw probabilities on TCGA-LAML (mean 0.14), and saturates again on the 39-AML in-house cohort and the n = 10 in-house survival cohort. These behaviours are part of the same observation: the threshold τ = 0.31 chosen on the pooled training corpus does not transfer linearly across out-of-distribution bulk and single-cell datasets. The signal that survives across cohorts without relying on threshold-based classification is the BeatAML drug-AUC correlation. The model is research-use only and has not been validated, certified, or approved for clinical, diagnostic, or therapeutic decision-making.

## Discussion

### Summary of what the data show

The Feature-Token Transformer described here was trained on a pooled scRNA-seq corpus (six samples from GSE123902 lung adenocarcinoma plus three primary AML samples; 32,342 cells, 13,369 genes) and produced strong discrimination of the training-class labels under cell-level stratified 5-fold cross-validation (mean fold AUROC 0.94) and a held-out test split (AUROC 0.94, MCC 0.70). The moderate MCC partly reflects label noise from the cross-cancer positive class and our deliberate avoidance of cross-corpus batch correction, which preserves deployability on unseen cohorts. It is more likely improved by additional within-AML training data than by further tuning of the current corpus. Knowledge distillation to a 1,000-gene student model preserved this discrimination at >85% class agreement and ρ ≈ 0.96 score correlation. Applied without retraining or threshold re-tuning to five external/application cohorts, the score behaved differently in each, and we report those differences openly.

The clearest external signal was the association between the predicted score and ex-vivo drug response in BeatAML. Across 452 patients and seven AML-relevant drugs spanning induction-chemotherapy backbones, the BCL-2 inhibitor venetoclax, and FLT3 inhibitors, per-drug Spearman correlations were uniformly positive in the 0.41-0.53 range. The aggregate correlation across the patient × drug matrix reached r = +0.482 (n = 3,164 patient-drug pairs); we treat the per-drug consistency as primary evidence, since pairs from the same patient are not fully independent. Mean predicted probability had substantial spread across the unit interval, in contrast to the saturation observed on out-of-distribution bulk cohorts. Survival outcomes and drug-response measurements were not used during model training, threshold selection, or model tuning, so the BeatAML drug-AUC association reflects external rank-order transfer rather than a supervised property of the model.

Drug-resistance can hamper treatments and reduce overall survival, but on the other hand, initial responses to treatments are often deep and resistance emerges only later. For this reason, we decided not to establish prior assumptions about correlation to overall survival but nevertheless evaluated it. The score did not stratify overall survival in TCGA-LAML at the available sample size (best log-rank p = 0.14, Cox HR 0.73 with p = 0.39 across n = 149 with usable survival metadata). It also did not stratify survival in our in-house n = 10 scRNA-seq cohort, in which predicted high-score fractions saturated at 81.6-96.8% and offered no within-cohort variance against which to test. Thus, we do not present the score as a survival-prediction tool; the BeatAML drug-AUC association is the result that should be cited from this work.

### Calibration and the transfer problem

The result that requires honest discussion is the behaviour of the fixed threshold τ = 0.31 across cohorts with different processing pipelines. On GSE74246 (sorted hematopoietic populations, bulk RNA-seq), the threshold called 79 of 81 samples persister, with the per-cell-type mean probability at or near 1.0 across stem, progenitor, blast, T-cell, NK-cell, monocyte, and erythroid populations; only B-cells showed variability (mean 0.35). This is not the stemness gradient that would have been the cleanest possible biological validation. We do not present it as such, and we do not include any earlier visualisation that suggested a gradient. The GSE74246 saturation reflects a library-type/scale mismatch rather than biology: after CPM-log1p the model genes sit systematically high relative to the droplet-scRNA-seq training distribution, so absolute scores saturate while within-cohort ranking is preserved reinforcing the recommendation to apply the score as a cohort-relative ranker.

On TCGA-LAML, the same trained model produced raw probabilities concentrated near zero (mean 0.14, median 4.4 × 10⁻⁵), and the prespecified threshold produced a degenerate stratification. Rather than discard the cohort, we report TCGA-LAML stratification using empirical percentile cuts and document this as an exploratory deviation from the prespecified analysis protocol. None of the percentile-based splits reached statistical significance for survival.

These two observations-saturation at the high end on GSE74246 and at the low end on TCGA-LAML - indicate that the threshold τ = 0.31, selected on the pooled training corpus, does not transfer cleanly to bulk RNA-seq cohorts processed under different pipelines.

These results suggest that the absolute probability scale is cohort-dependent, whereas the within-cohort rank ordering remains informative. This is supported by the BeatAML drug-AUC association, which depends only on rank ordering, not on the threshold. We therefore recommend that future users apply the score as a relative ranker on each new cohort (e.g. score quantiles or rank-based comparisons) rather than as an absolute classifier at any fixed cutoff. If a hard cutoff is needed, cohort-specific recalibration (for example, isotonic regression on a held-out portion of each new cohort) should precede threshold-based classification. The threshold-based prevalence numbers reported here (97.5% on GSE74246, 50% under percentile-top-50 on TCGA-LAML, 83.8% mean on the 39-AML cohort) should be read as descriptions of model behaviour rather than as biological prevalence estimates.

### What the score appears to capture

Three orthogonal pieces of evidence support the interpretation that the rank ordering of the score reflects biology related to drug-tolerance and stemness, even though the threshold calibration is unreliable: the BeatAML drug-AUC correlation, consistent across seven drugs and 3,164 patient-drug pairs from 452 patients; the enrichment of the 1,000-gene panel for AML genetic dependencies (DepMap mean gene-effect −0.42 vs. background −0.10, p < 0.001 by permutation); and the enrichment for EMT, adhesion, Wnt/β-catenin, Hippo signalling, and transmembrane receptor tyrosine kinase activity (including FLT3) - programs that have been described in drug-tolerant states across cancer types[1,2,18]. These are supportive observations, not definitive identification of a persister phenotype, and we discuss them as such.

What we did not do is train on a within-AML labelled persister/non-persister contrast, because no such densely-labelled scRNA-seq dataset was available to us. The training corpus uses lung-cancer metastasis vs. lung-normal/primary as one component of the positive/negative contrast, with three AML samples added to the positive class. The cross-cancer assumption-that a state learned partly from lung adenocarcinoma is informative for AML-rests on the fact that the pathways enriched in our 1,000-gene panel (epithelial-mesenchymal transition, Wnt/β-catenin, Hippo signalling, focal adhesion, transmembrane RTK activity) are programs reported as conserved features of drug-tolerant states across multiple cancer types[1,2,18], rather than lineage-specific features of any one tumour origin. We therefore interpret the score as capturing a transferable transcriptional component of treatment-relevant malignant heterogeneity, and we treat the BeatAML drug-AUC association as the empirical anchor that this interpretation is at least not fatally wrong. The label “transcriptomic score” is a working interpretation; “transcriptomic score that overlaps with persister biology” is the more cautious reading and is equally consistent with the data. A future revision will be trained on a within-AML treatment-naïve vs. post-treatment contrast as soon as such data become available.

Across independent cohorts the score behaved consistently with a conserved epithelial/stemness programme rather than a malignancy-or persister-specific signal. In sorted hematopoietic populations (GSE74246) it scored highest in primitive HSC/LSC/blast fractions and lowest in mature cells. In a lung adenocarcinoma atlas (GSE131907), per-compartment scores tracked malignant-epithelial content-high in pleural-effusion, bronchoscopy and metastatic compartments, and indistinguishable between primary tumour and normal lung, consistent with the low malignant-cell fraction of dissociated primary tumour samples. In a colorectal cohort (GSE200997), the score did not separate tumour from normal at the sample level (16 vs 7 patients, p = 0.41); a simpler mean-expression baseline behaved identically, indicating this is a property of the signal rather than the architecture. We therefore interpret the score as a within-cohort relative ranker of stemness/tolerance-associated transcription, not a tumour-versus-normal classifier -a boundary we state explicitly so the score is applied appropriately.

To assess whether the persister-score signal reflects canonical biological ageing rather than a therapy-stress-specific programme, we projected conserved cross-species transcriptomic ageing signatures derived from a recent multi-tissue, multi-species integration study onto AraC-treated AML blast cells from a published single-cell dataset (GSE146590; Duy et al.). Both the conserved ageing-up and ageing-down gene sets were elevated simultaneously in therapy exposed malignant cells compared to untreated controls, a pattern inconsistent with directional biological ageing and more consistent with generalised transcriptional stress dysregulation. The composite biological age score increased modestly in AraC-treated cells (fold change 1.31; Mann-Whitney effect size r = 0.069, p = 0.001), whereas the SASP-stemness co-elevation signal that defines the hybrid state showed a substantially larger effect (r = 0.466 and r = 0.337 respectively). Notably, biological age score was negatively correlated with stemness specifically in therapy-exposed cells (Spearman r = −0.162, p = 1.1 × 10⁻⁸), a relationship absent in untreated controls (r = +0.006, p = 0.80), suggesting that stemness maintenance and canonical ageing programmes oppose one another under chemotherapy pressure. Hybrid state cells defined by co-elevation of senescence arrest, SASP, and stemness were not significantly more biologically aged than other malignant cells (Mann-Whitney p = 0.163). Together these observations indicate that the hybrid transcriptional state identified here is a therapy specific adaptation rather than a manifestation of accelerated cellular ageing, and that the persister score signal is grounded in stemness and stress-response biology rather than in the inflammaging programmes that characterise normal tissue ageing.

### Candidate target list and intended use

The 250-candidate ranked membrane-protein list, with its five-tier partitioning (SAFE n = 9, CHECK n = 11, CAUTION n = 19, HIGH_RISK n = 17, EXCLUDE n = 194), is a hypothesis-generating output for downstream functional work. The CAUTION tier recovered FLT3 and CD33 from the unbiased ranking - both established AML therapeutic targets with clinical precedent-providing a positive control that the prioritisation pipeline recovers biologically meaningful candidates. Individual candidate identities beyond publicly known AML targets are retained as proprietary pending IP review; a confidential reviewer supplement is available on request. No wet-lab validation has been performed; the intended use of the list is prioritisation in subsequent perturbation, antibody-screening, or chemical-probe experiments.

The 91.6% concordance between the transcriptomic ranking and HPA’s independent protein-level surface annotation strengthens the interpretation that the prioritisation pipeline is selecting for plausibly tractable surface targets, though true protein-level surface expression on primary AML blasts remains to be confirmed experimentally.

### Comparison with prior work

The present study contrasts with prior single-cell AML persister characterisation in three ways. First, where most prior work uses bulk transcriptomics with pathway-level inference, we provide cell-level scores with a published, frozen model. Second, where prior single-cell classifiers have often been evaluated on the same cohorts they were trained on, we report sample-level cross-validation, a held-out test split, and four independent application cohorts, with all transfer behaviours-including the saturation problems-documented openly. Third, where prior work has typically stopped at characterisation, we link the model output to genetic-dependency data (DepMap), pathway enrichment, and a normal-tissue-filtered candidate-target list. We do not claim better discrimination than prior work; with the calibration limitations described, the appropriate framing is “strong rank-ordering on independent drug-response data, with explicit mechanistic anchoring,” not “improved transcriptomic score classifier.”

## Limitations

We recognise the following limitations and document them so that downstream users of the score and reviewers of this paper understand the boundaries of the work.

- **Training-corpus composition.** The training data include lung-cancer-derived scRNA-seq as a component of the positive/negative contrast, with three primary AML samples in the positive class. The model has not been trained on a within-AML treatment-naïve vs. post-treatment contrast and is therefore best understood as a transcriptomic score with cross-tumour generalisation rather than as an AML-specific transcriptomic score classifier in the strict sense.
- **Aliquot-grouped cross-validation is not true donor-level LOGO.** Beyond the headline cell-level 5-fold StratifiedKFold, we performed aliquot-grouped 3-fold StratifiedGroupKFold validation as a leakage-reduction analysis (FIGURE 4B, Supplementary Table S4). However, several of our 9 sample groups originate from the same donor: ALIQUOT A1 and ALIQUOT A2 are aliquots of Donor A, and DONOR B (NORMAL LUNG) and DONOR B (PRIMARY TUMOUR) are normal-lung and primary-tumour samples from Donor B. The reported aliquot-grouped AUROC therefore reduces within-sample cell leakage relative to cell-level CV but is not equivalent to true leave-one-donor-out (LOGO) validation using patient identifiers. True donor-level LOGO using independent patient identifiers remains future work.
- **Calibration does not transfer cleanly across bulk cohorts.** The fixed threshold τ = 0.31 saturates at the high end on GSE74246 (97.5% positive) and at the low end on TCGA-LAML (mean probability 0.14). Threshold-based prevalence numbers should be interpreted as model-behaviour descriptions rather than biological prevalence estimates. A future revision would benefit from cohort-specific recalibration (for example, isotonic regression on a held-out portion of each new cohort) before threshold-based classification.
- **TCGA stratification was exploratory.** Because the prespecified threshold was non-informative on TCGA-LAML, we reported survival stratification using empirical percentile cuts. This is an exploratory deviation from the prespecified protocol. We tested multiple stratification methods and report all of them; we did not apply formal multiple-testing correction across stratification methods. The cohort showed no significant survival association under any method.
- **The in-house n = 10 scRNA-seq cohort is uninformative for survival.** Predicted high-score fractions saturated at 81.6-96.8% across all ten patients. The cohort is too small and too saturated to test a survival association meaningfully and is reported as such.
- **No within-AML negative control for the 39-donor cohort.** We do not have an independent set of AML samples enriched for non-persister states; we therefore cannot determine whether the high persister-call rate (mean 83.8%) on the 39 in-house donors reflects real AML-persister enrichment or model behaviour on AML transcriptomes generally.
- **The B-cell exception in GSE74246 is unexplained.** B-cells were the only cell type with substantial variability in predicted probability. We have not formally identified the transcriptional features driving this and do not claim it as a meaningful biological signal. The most parsimonious explanation is the absence of B-cell representation in the training corpus combined with the lymphoid-vs-myeloid transcriptional distance: the pooled training corpus is dominated by epithelial (lung) and myeloid (AML) lineages, so mature B-cells sit further from any learned class boundary than the other GSE74246 populations and the score is correspondingly less saturated. We flag this as a behaviour rather than a finding, and would expect retraining on a corpus that includes lymphoid cells to either confirm the explanation or expose a different one.
- **Survival in BeatAML is not significant.** The Cox HR for the score was 1.22 (p = 0.19); the strongest stratification (extreme quartiles) gave log-rank p = 0.16. We do not claim BeatAML survival prediction. The drug-AUC correlation, not survival, is the BeatAML result.
- **Architectural contribution is regularisation-driven transferability, not best-in-class internal performance.** On internal cross-validation, simpler classifiers (L2-regularised logistic regression, random forest) achieve higher AUROC than the Transformer on the training task. The Transformer’s contribution is that its regularisation produces a score that transfers to the BeatAML drug-AUC anchor (aggregate Spearman r = +0.482), where the same simpler classifiers fail to transfer (LR aggregate r = +0.167, RF r = −0.053). This pattern is consistent with the Transformer learning a smoother, less donor-specific decision function. We report this honestly rather than framing the architecture as the best-on-training method.
- **The candidate-target list is not experimentally validated.** Wet-lab confirmation of any candidate has not been performed. The list is intended as a starting point for follow-up experimental work, not as a list of validated therapeutic targets.
- **Research-use only.** The model is not validated for clinical decision-making. We make no clinical claims.

## Conclusions

We trained a Feature-Token Transformer on a pooled scRNA-seq corpus and applied it to four independent cohorts without retraining or threshold re-tuning. The model discriminates the training-class labels strongly under cross-validation and on a held-out test split (mean fold AUROC 0.94, held-out test AUROC 0.94). Distillation to a 1,000-gene student preserves performance at >85% class agreement.

The principal external result is the association of the predicted score with ex-vivo drug-response AUC in BeatAML across seven AML-relevant drugs (per-drug Spearman r = 0.41-0.53, all p < 10⁻²¹; aggregate r = +0.482 across 3,164 patient-drug pairs from 452 patients, with pairs from the same patient not fully independent). In a large independent AML cohort, higher predicted score is associated with reduced ex-vivo drug responsiveness across induction-chemotherapy backbones, BCL-2 inhibition, and FLT3 inhibitors. Survival and drug-response data were not used in training, threshold selection, or model tuning, so this association reflects external rank-order transfer rather than a supervised property of the model.

Threshold-based classification did not transfer cleanly across bulk cohorts. We report the GSE74246 saturation, the TCGA-LAML score-distribution shift, and the in-house cohort saturation transparently. We interpret these limitations as reflecting unreliable absolute calibration but informative rank ordering, an interpretation supported by the BeatAML drug-response correlation. The score does not stratify overall survival in TCGA-LAML or in our small in-house scRNA-seq cohort, and we do not propose it as a survival-prediction tool.

Mechanistic anchoring through DepMap essentiality and KEGG / GO / MSigDB Hallmark / WikiPathways enrichment supports the interpretation that the score is grounded in plausible drug-tolerance-relevant biology (EMT, adhesion, Wnt/β-catenin, Hippo, transmembrane RTKs including FLT3). The 250-candidate ranked surface-protein list, with FLT3 and CD33 recovered as clinically-validated positive controls in the CAUTION tier, is provided as a starting point for follow-up functional work.

In summary, we present a Transformer-derived transcriptomic score that shows external rank-order association with ex-vivo AML drug response in BeatAML. The score did not stratify overall survival, and its fixed threshold did not transfer reliably across all external cohorts. These findings support the model as a research-use ranking tool for drug-response-associated transcriptional states in AML, not as a clinical classifier. Future work should focus on donor-level validation, cohort-specific calibration, training on within-AML treatment-response data, and experimental testing of candidate targets.

## List of abbreviations

AML: Acute myeloid leukaemia
scRNA-seq: Single-cell RNA sequencing
AUROC: Area under the receiver operating characteristic curve
AUPRC: Area under the precision-recall curve
MCC: Matthews correlation coefficient
CV: Cross-validation
LOGO: Leave-one-donor-out
PCA: Principal component analysis
CPM: Counts per million
DE: Differential expression
FDR: False discovery rate
DTP: Drug-tolerant persister
LSC: Leukaemic stem cell
HSC: Haematopoietic stem cell
DepMap: Cancer Dependency Map
TCGA: The Cancer Genome Atlas
LAML: Acute Myeloid Leukaemia (TCGA cohort)
CRC: Colorectal cancer
AUC: Area under the curve
TKI: Tyrosine kinase inhibitor
CMS: Consensus molecular subtype
HGNC: HUGO Gene Nomenclature Committee.

## Declarations

### Ethics approval and consent to participate

Use of de-identified scRNA-seq samples from the in-house University of Helsinki AML cohort was approved under the institutional ethical review covering the FIMM AML translational research programme. All participants provided written informed consent for research use of their samples in accordance with the Declaration of Helsinki. Public datasets (GSE123902, GSE74246, BeatAML, TCGA-LAML, DepMap, HPA, GTEx) were used under the data-access terms of their respective providers.

### Consent for publication

Not applicable. No identifiable individual-participant data are included in this manuscript or supplementary materials.

### Availability of data and materials

Public datasets used in this work are available through their original sources: GSE123902 and GSE74246 (NCBI GEO); BeatAML (https://biodev.github.io/BeatAML2/); TCGA-LAML (NIH Genomic Data Commons); DepMap Public 24Q2 (https://depmap.org/portal/); Human Protein Atlas (https://www.proteinatlas.org/); GTEx Portal (https://www.gtexportal.org/).

### Author contributions

JB conceived the study, designed and implemented the computational pipeline, performed the analyses, and drafted the manuscript(Writing-original draft; Writing-review & editing). CH contributed to study design, supervised the project, and edited the manuscript(Supervision; Writing-review & editing). MV-K contributed to study design, provided scientific input on AML biology and ex-vivo drug-response data, supervised the project, and edited the manuscript. (Supervision; Writing-review & editing) SA assembled and curated the single-cell and clinical sample data, compiled the cohort characteristics, and contributed to interpretation of the scRNA-seq samples (Data curation; Writing-review & editing) All authors read and approved the final manuscript.

Single-cell RNA-seq samples from the in-house University of Helsinki AML cohort (used as three positive-class samples in training and as the 39-donor independent application cohort and the n = 10 survival cohort) are subject to ethical and consent restrictions and are available from the authors upon reasonable request, conditional on appropriate data-access agreements.

## Acknowledgements

The authors thank the Finnish Hematology Registry and Clinical Biobank (FHRB) for access to patient samples and associated clinical data, and the FIMM single-cell sequencing unit / FIMM Technology Centre for sample processing and sequencing. We are grateful to the patients and healthy donors who consented to the use of their samples in research. We acknowledge CSC-IT Center for Science, Finland, for computational resources provided through an academic project allocation. During the preparation of this manuscript, the author(s) used Anthropic’s Claude (claude.ai) for language editing, suggestions on manuscript structure, and assistance with formatting of supplementary tables. No scientific content, analyses, results, or interpretations were generated by the AI tool. The author(s) reviewed and edited the content as needed and take full responsibility for the content of the publication.

## Code availability

Source code, trained model weights, scaler/PCA artefacts, the prespecified threshold, training gene order, training and inference logs, and the full reproducibility package are available at https://github.com/jyotidiplearning99/AML_Persister_Analysis. Supplementary Tables S1-S8 are provided with this manuscript. Additional candidate identities can be provided to editors and reviewers as confidential supplementary material where permitted by institutional IP review.

## Supplementary materials index

The following Supplementary Tables accompany this manuscript. They are provided as separate files (CSV/XLSX bundle).

**Supplementary Table S1.** Per-fold cross-validation metrics and held-out test confusion matrix. Stratified 5-fold AUROC (0.944, 0.945, 0.944, 0.908, 0.942; mean 0.936 +/- 0.014); selected threshold τ = 0.310; held-out test AUROC 0.941, MCC 0.696, F1 0.895, balanced accuracy 0.856; confusion matrix TN 1,763, FP 339, FN 552, TP 3,815.

**Supplementary Table S2.** Aliquot-grouped cross-validation performance for all five models across three folds (Transformer, logistic regression, random forest, LSC17, mean-expression).

**Supplementary Table S3.** BeatAML per-drug Spearman correlations with the predicted score (Venetoclax r = 0.47, Cytarabine 0.48, Daunorubicin 0.50, Idarubicin 0.52, Midostaurin 0.41, Gilteritinib 0.53, Quizartinib 0.52) and the aggregate correlation across 3,164 patient-drug pairs from 452 patients (r = +0.482; the aggregate p-value computed under independence assumptions is < 10⁻¹⁸⁴, but is inflated because pairs from the same patient are not fully independent-per-drug correlations are the primary evidence). Also includes the full TCGA-LAML survival stratification table across median, tertile, quartile, extreme-quartile, threshold, and Cox proportional-hazards methods.

**Supplementary Table S4**. Baseline classifier comparison against the Transformer on BeatAML drug-AUC. Includes aggregate Spearman correlations across 3,164 patient-drug pairs (all models on 1k and 13k panels) and per-drug Spearman correlations across the seven AML-relevant agents (Venetoclax, Cytarabine, Daunorubicin, Idarubicin, Midostaurin, Gilteritinib, Quizartinib) for the Transformer and four baselines (logistic regression, random forest, LSC17, mean-expression).

**Supplementary Table S5.** Per-cohort predictions across all five independent application cohorts (39 AML, GSE74246, TCGA-LAML, BeatAML, in-house n = 10 survival), provided as a multi-sheet Excel file. Includes per-sample mean and SD probability, persister-call counts at τ = 0.31, gene coverage, and (for GSE74246) cell-type and group annotations.

**Supplementary Table S6.** Full pathway-enrichment outputs for the 1,000-gene panel against KEGG, GO Biological Process / Molecular Function / Cellular Component, MSigDB Hallmark, WikiPathways, and DSigDB drug-perturbation libraries. Combined-score and adjusted-p columns provided. Top hits include Adherens junction, ECM-receptor interaction, Hippo signaling, Focal adhesion, Wnt signaling, EMT (Hallmark), Estrogen Response, p53 Pathway, KRAS Signaling, TGF-β signalling, Hypoxia, transmembrane receptor tyrosine kinase activity, and Wnt/β-catenin Signaling Pathway in Leukemia (WP3658). Pathway enrichment only; no transcription-factor claims are made in this manuscript.

**Supplementary Table S7.** Top-30 highest-priority candidates (9 SAFE + 11 CHECK + 10 highest-priority CAUTION) with full annotation: priority score, safety tier, mechanism class (Membrane / GPCR / Ion channel / Transporter / RTK / CD marker), therapeutic approach (Antibody / Small molecule / Clinical precedent / TKI / TBD), HPA high-and medium-expression tissue counts, GTEx maximum critical-tissue expression, GTEx hematopoietic-tissue expression, hematopoietic-tissue detection flag, and the safety-rank ordinal. HPA protein-level concordance columns are also provided: HPA_surface_protein flag (Yes/No), HPA_antibody_reliability tier (Enhanced / Supported / Approved / Uncertain / Not available), subcellular location, and protein class. The full HPA tissue-by-tissue expression matrix for these 30 candidates is provided in the confidential reviewer supplement.

**Supplementary Table S8.** Full 250-candidate ranking with composite priority score, tier assignment, and per-feature contributions (HPA high/medium-tissue counts, GTEx critical-tissue expression, GTEx hematopoietic-tissue expression, ubiquitin/CD/JAK-STAT/BCL2/MYC flags, therapeutic-approach class, adjusted priority). Tiers: SAFE (n = 9), CHECK (n = 11), CAUTION (n = 19; includes FLT3 and CD33 as clinically-validated positive controls), HIGH_RISK (n = 17), EXCLUDE (n = 194). HPA protein-level concordance columns are also provided per gene: HPA_surface_protein flag, HPA_antibody_reliability tier, subcellular location, and protein class. 248 of 250 candidates (99.2%) matched HPA on HGNC symbol; the two unmatched (C16ORF74, C19ORF33) use older nomenclature and were resolved manually.

### Note on transcription-factor regulon analysis

A DoRothEA-based TF regulon analysis was originally planned but is reported separately in a forthcoming companion analysis. The present manuscript does not include TF claims and Supplementary Table S6 contains pathway-level enrichment only.

### Supplementary File

Confidential reviewer supplement with unanonymised candidate-target identities is available to editors and reviewers on request.

